# Distinct Conformations of Mirabegron Determined by MicroED

**DOI:** 10.1101/2023.06.28.546957

**Authors:** Jieye Lin, Johan Unge, Tamir Gonen

## Abstract

Mirabegron, commonly known as “Myrbetriq”, has been widely prescribed as a medicine for overactive bladder syndrome for over a decade. However, the structure of the drug and what conformational changes it may undergo upon binding its receptor remain unknown. In this study, we employed microcrystal electron diffraction (MicroED) to reveal its elusive three-dimensional (3D) structure. We find that the drug adopts two distinct conformational states (conformers) within the asymmetric unit. Analysis of hydrogen bonding and packing demonstrated that the hydrophilic groups were embedded within the crystal lattice, resulting in a hydrophobic surface and low water solubility. Structural comparison revealed the presence of *trans*- and *cis*-forms in conformers **1** and **2**, respectively. Comparison of the structures of Mirabegron alone with that of the drug bound to its receptor,^1^ the beta 3 adrenergic receptor (β3AR) suggests that the drug undergoes major conformational change to fit in the receptor agonist binding site. This research highlights the efficacy of MicroED in determining the unknown and polymorphic structures of active pharmaceutical ingredients (APIs) directly from powders.

---

Mirabegron, marketed as “Myrbetriq”, is the first beta3-adrenergic (β3AR) agonist available for clinical use in the treatment of overactive bladder syndrome.^2,3^ Its chemical structure comprises various groups, including phenylethanolamine, acetanilide, and 2-aminothiazole, connected by aliphatic chains (Figure 1A). The molecule as a whole is hydrophobic and exhibits poor water solubility,^4^ despite containing hydrophilic hydroxyl, amine, and carbonyl groups. Understanding the 3D crystal structure of Mirabegron is essential for comprehending its properties and formulation. Despite its wide prescription and ranking as one of the top 160 prescribed medicines in the United States in 2020, with over 3 million prescriptions,^5,6^ the crystal structure of Mirabegron remained unknown. Mendoza *et al*. reported the PXRD spectrum of Mirabegron, identifying it as a triclinic space group P1 (a=5.35Å, b=11.61Å, c=17.59Å, α=70.79°, β=84.42°, γ=86.29°).^7^ Tamayo *et al*. later reported PXRD and DFT refinement for Mirabegron, but without an available structure.^8^ In a single particle Cryo-EM study of Mirabegron bound to dog β3AR, Mirabegron adopted a conformation that was different than the expected conformation based on the synthesis alone. Although the experimental density for Mirabegron in that study was poor the structure indicated that the phenylethanolamine moiety binds to the highly conserved orthosteric site of β3AR, while the 2-aminothiazole ring interacts with the receptors’ extracellular face (exosite) when interacting with its receptor (PDB entry: 7DH5).^1^

**Figure 1.**
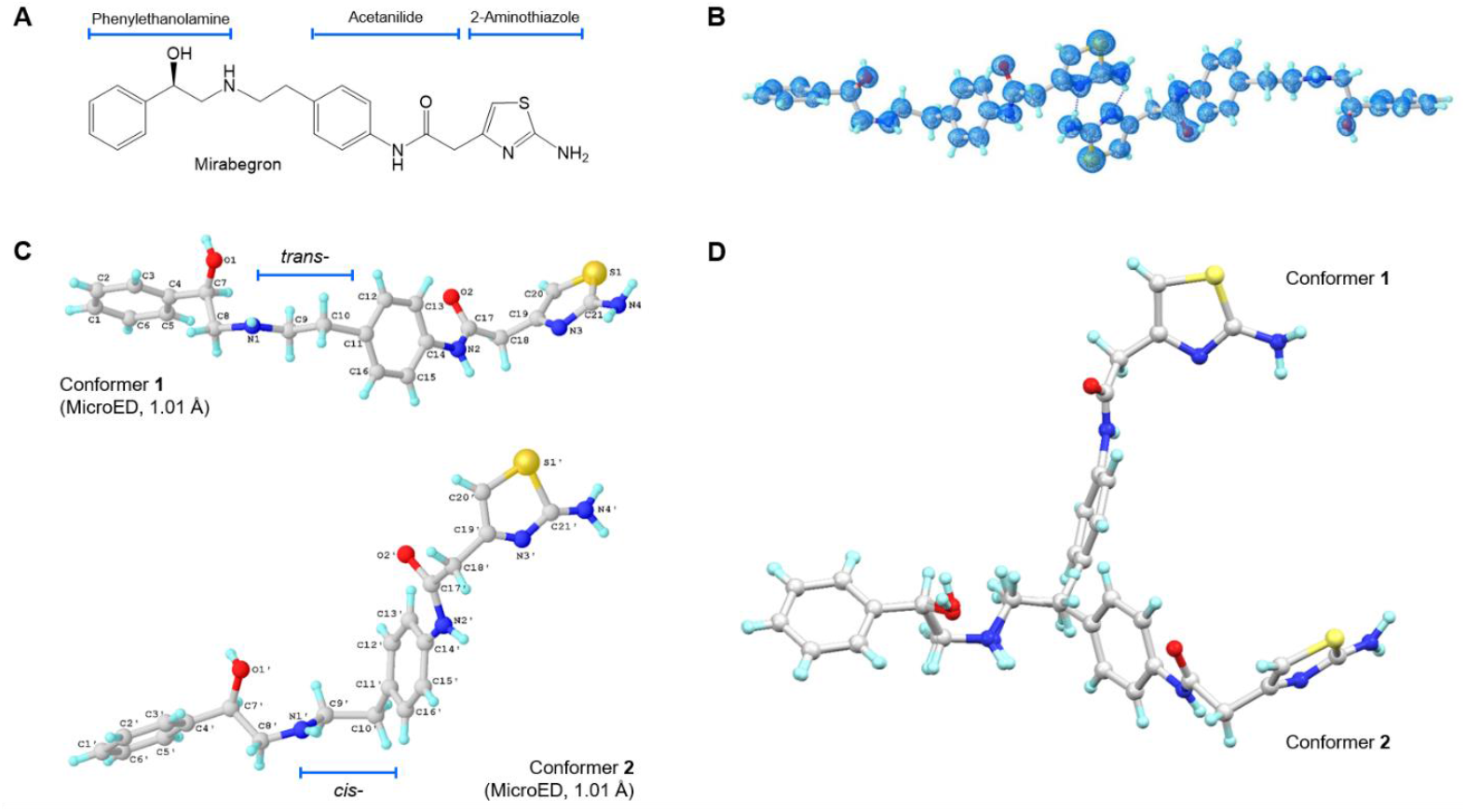
MicroED structures of Mirabegron: (A) Chemical structure; (B) MicroED structure of Mirabegron at 1 Å resolution. The 2F_o_-F_c_ map is shown in blue; (C) Conformers **1** and **2** (*trans*- and *cis*-, respectively) observed in the unit cell; MicroED structure of Mirabegron was solved by SHELXT^28^ and refined using SHELXL^29^. (D) Overlay of the two conformers.

The main challenge in conventional single crystal X-ray diffraction (SC-XRD) is the difficulty in obtaining large crystals from powdery substances. For synchrotron SC-XRD, crystals of at least 5 μm are typically required.^9^ This poses a challenge for small molecules like Mirabegron, which have a linear structure with several single bonds, allowing flexibility in conformation with minimal steric hindrance and rotational barriers in the aliphatic chains.^10^ However, Microcrystal Electron Diffraction (MicroED) has emerged as a complementary technique to conventional crystallography methods.^11,12^ It has proven useful in analyzing crystals that are just a billionth the size needed for SC-XRD and for small molecules directly from seemingly amorphous powders that often contain nanocrystals. This is particularly advantageous for the active pharmaceutical ingredients (APIs) development or novel drug structures. For example, MicroED was utilized to solve elusive crystal structures of drugs like Bismuth subgallate,^13^ Methylene blue derivative (MBBF4),^14^ Orthocetamol,^15^ Lomaiviticins,^16^ Levocetirizine dihydrochloride.^17^ Another application is the screening of different polymorphs, salts, co-crystals, and solvates of drug molecules, such as the polymorphs of Remdesivir,^18^ Indomethacin,^19^ Vemurafenib;^20^ co-crystal structure of 2-Aminopyrimidine succinic acid (2:1);^21^ Olanzapine salts;^22^ and Methyl piperazine-1-carboxylate as a solvated crystal.^23^ Here we use MicroED to determine the atomic structure of Mirabegron that has remained elusive for decades despite its wide use and FDA approval.^5,6^

The sample preparation for MicroED followed the previously described procedure (details can be found in the Supporting Information).^24^ The grid containing the crystals was examined using the Thermo Fisher Talos Arctica Cryo-TEM operating at 200 kV and approximately 0.0251 Å wavelength. The microscope was equipped with a CetaD CMOS camera and EPUD software. The crystal thickness played a crucial role in obtaining optimal diffraction, so crystals were initially screened using imaging mode (LM 210×) using a grid atlas. Only thin plate crystals with lighter contrast were selected for further analysis (see Figure S1 in the Supporting Information). The eucentric heights of the crystals were manually calibrated (SA 3400×) to ensure proper centering during the continuous rotation. For data collection, a 70 μm C2 aperture and a 50 μm selected area (SA) aperture were utilized to reduce background noise and achieve a 1.4 μm beam size. The typical data collection involved a constant rotation rate of approximately 1 °/s over an angular range of 100° (−50° to +50°), with an exposure time of 1 second per frame. The MicroED movies were converted from mrc format to smv format using the mrc2smv software (available freely at https://cryoem.ucla.edu/microed).^25^ High-quality datasets were indexed and integrated using XDS,^26,27^ resulting in a completeness of over 50% for each dataset (see Table S2 in the Supporting Information) which increased to 99.7% after scaling and merging data from 3 individual crystals (see Table S1 in the Supporting Information). The intensities were converted to SHELX hkl format using XDSCONV.^27^ The MicroED structure was solved *ab initio* using SHELXT^28^ at a resolution of 1.01 Å and subsequently refined with SHELXL^29^ to achieve a final R_1_ value of 16.6% (see Figure 1 and Table S1 in the Supporting Information). The positions of heavier atoms were accurately determined from the charge density map (see Figure 1B). Since not all hydrogen (H) atoms could be located at this resolution, their positions were refined using a combination of constrained and free approaches.

The MicroED structure of Mirabegron was determined to be a non-centrosymmetric triclinic space group P1, with unit cell dimensions of a=5.27 Å, b=11.58 Å, c=17.27 Å, α=70.73°, β=84.35°, and γ=86.37°. In the asymmetric unit, two distinct conformers, referred to as “conformer **1**” and “conformer **2**,” were identified (Figure 1C; Notations see Scheme S1 in Supporting Information). The simulated powder X-ray diffraction (PXRD) spectrum matched well with the previously reported PXRD data (Figure S2 in Supporting Information).^7^ Within the MicroED structure, conformer **1** and conformer **2** form tightly stacked pairs through eight hydrogen bonds, creating a dense three-dimensional network (Figure 2; Table S3 in Supporting Information). The hydrogen bonds can be categorized into two groups: (1) repetitive O1‒H···N1/O1’‒H···N1’ and N2‒ H···O2/N2’‒H···O2’ along the *a*-axis. The former exhibits shorter distances (2.8-2.9 Å) compared to the slightly longer distances observed in the latter (3.0-3.3 Å). Both types of hydrogen bonds contribute to the tight packing along the *a*-axis (Figure 2). (2) Interactive N4‒H···N3’/N4’‒H4···N3 hydrogen bonds occur between the 2-aminothiazole rings. These bonds have longer distances (3.1-3.2 Å), but the additional N4‒H···O2/N4’‒H···O2’ (2.9-3.1 Å) interactions with the carbonyl group help establish a complex hydrogen bonding network involving N2’, O2, N4, N3, N2, O2, N4’, N3’, and adjacent carbon atoms. The *zig-zagged* hydrogen bonding network between crystal layers stabilizes and extends the packing along the *b* and *c* axes (Figure 2). The space-filling packing diagram of Mirabegron reveals a dense packing arrangement in the lattice. This is achieved through pairs of hydrogen bonds and *van der Waals* contacts such as C‒H···π or N‒ H···π interactions.^30^ The hydrophilic hydroxyl, amine, and carbonyl groups are embedded within these pairs, resulting in a hydrophobic surface and low water solubility (Figure S3 in Supporting Information). For more detailed information, please refer to the Supporting Information.

**Figure 2.**
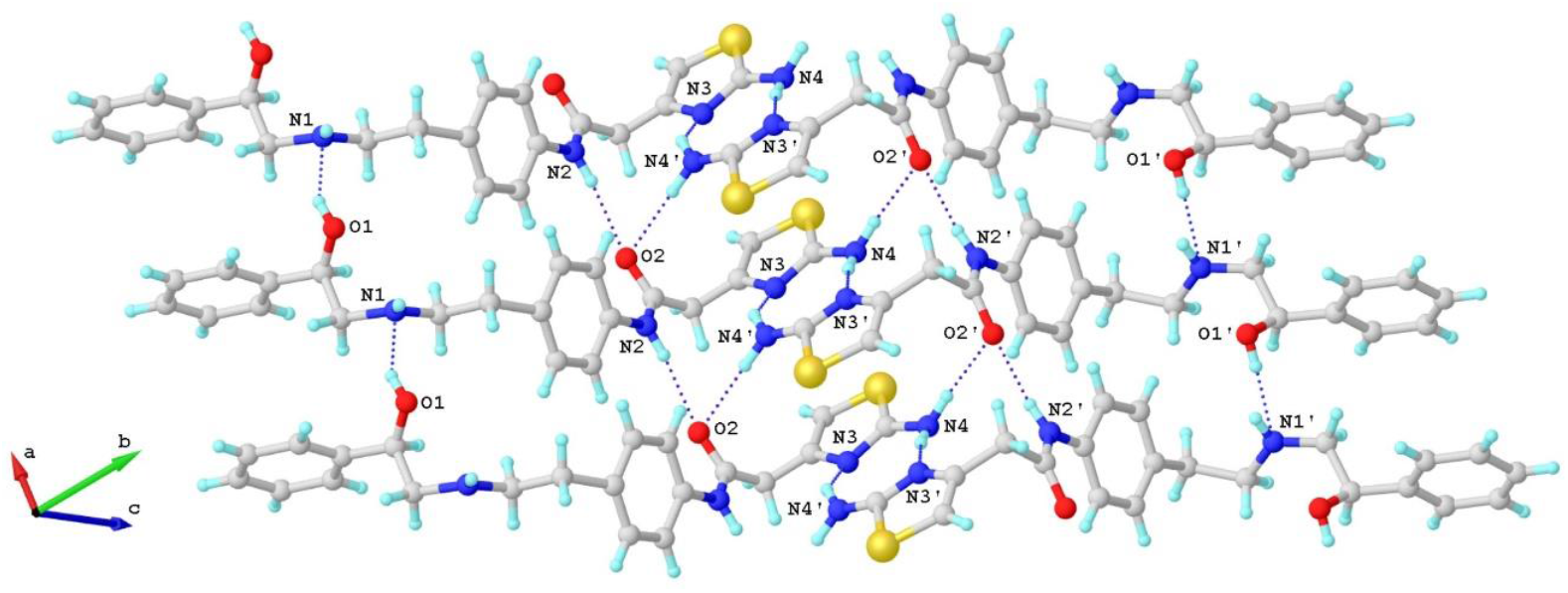
Hydrogen-bond interactions in Mirabegron crystal packing. The hydrogen-bond interactions are represented by the dashed lines. The contact atoms are labeled.

In solution, the C‒C and C‒N bonds of Mirabegron have the ability to rotate freely, allowing for flexible conformations. However, in the crystal lattice, two distinct conformers, labeled as conformer **1** and conformer **2**, were observed. These conformers formed pairs and were arranged in a “head-to-tail” fashion within the crystal lattice. It is important to note that the structural parameters of these conformers, including bond lengths, bond angles, and torsion angles, exhibit noticeable variations depending on the specific chemical environment. Upon comparing conformer **1** and conformer **2**, it was observed that most bond lengths within the aliphatic chains displayed very small differences, typically less than 0.2 Å. Furthermore, the majority of carbon and nitrogen atoms maintained an sp^3^ geometry, resembling a pyramidal shape, resulting in bond angles close to 109.5°. Experimental analysis revealed that the two aromatic 2-aminothiazole rings in Mirabegron were non-identical and exhibited slight distortions. These distortions were likely influenced by different atom displacements and hydrogen bond interactions present in diverse chemical environments.^31^

Comparing conformer **1** and conformer **2**, significant differences were observed in torsion angles, which have a greater impact on the overall conformation than bond lengths or angles (Figure S4 in Supporting Information). The primary differences include: 1. Rotation of C9‒C10/C9’‒C10’ bonds, resulting in *trans*- and *cis*-forms of the phenylethanolamine and acetanilide groups. Despite the higher energy and structural hindrance of the *cis*-form, hydrogen-bond interactions compensate for the rotation barrier, enabling the coexistence of *trans*- and *cis*-conformers that facilitate “head-to-tail” packing. 2. Rotation of C14‒N2/C14’‒N2’ bonds controlled by antiparallel orientations of N2‒H···O2 and N2’‒H···O2’ hydrogen bonds. This leads to approximately 30° angles between C17/C17’ and the phenyl rings, resulting in a staggered arrangement. 3. Rotation of C17‒C18/C17’‒C18’ and C18‒C19/C18’‒C19’ bonds maximizes hydrogen bond interactions (N4‒H···N3’/N4’‒H···N3, N4‒H···O2/N4’‒H···O2’) between the two 2-aminothiazole rings. These interactions tightly bind the crystal layers along the *b* and *c* axes in an arched direction. These differences in torsion angles play a crucial role in determining the conformation and packing arrangement of the molecule.

We next compared the structure of Mirabegron on its own with the previously determined structure of the drug bound to its receptor. The structure of the Mirabegron-dog β3AR complex was determined by single particle Cryo-EM analysis at a resolution of 3.16 Å (PDB entry: 7DH5) (Figure 3).^1^ The drug was found buried deep in the agonist binding pocket of the receptor on the extracellular side of the membrane. Several amino acids formed hydrogen bonds with specific regions of Mirabegron. For instance, Asn341 and Asp126 at the orthosteric site formed hydrogen bonds with O1 and N1, respectively. Similarly, Phe207 and Cys205 at the extracellular site (exosite) formed hydrogen bonds with N2 and N4 of Mirabegron (Figure S7 in Supporting Information). In the crystal lattice, these interactions were replaced by hydrogen bond interactions involving O1···N1/O1’···N1’, N2···O2/N2’···O2’, N4···O2’/N3’, and N4’···O2/N3’ (Figure 2, Table S3 in Supporting Information). When comparing conformer **1** and conformer **2** of Mirabegron with the receptor bound structure it was evident that neither conformer could directly enter the protein agonist binding site without significant conformational change in the drug itself (Figure 3). In order for the phenylethanolamine side of Mirabegron to interact with the highly conserved orthosteric sites of the receptor, the C8‒N1 bonds would have to rotate, resulting in the rearrangement of C7 and C9 from *cis*-to *trans*-form (Figure S5 in Supporting Information). This rotation alone would allow conformer **1** to enter the agonist binding pocket of β3AR although conformer **2** would require additional conformational changes. Conformer **2** would also require the rotation of both the C8’‒ N1’ and C9’‒C10’ bonds to flip almost half of its structure into the *trans*-form (Figure S6 in Supporting Information). We note that conformer **2** fits nicely in an allosteric site proximal to the putative binding pocket. The 2-aminothiazole side of Mirabegron interacted with three amino acids at the exosite, which is partially outside the agonist binding site in β3AR (Figure 3). This interaction involved hydrogen bonds between N2/N2’ and N4/N4’ of Mirabegron with the tripeptide Cys205‒ Ala206‒Phe207, where N2/N2’ and N4/N4’ served as donors that hydrogen bonded to the carbonyl group of Cys205 and Phe207 with distances of 3.1 Å and 3.5 Å, respectively (Figure S7 in Supporting Information). Unlike the orthosteric site, the exosite allowed for more flexibility, requiring additional bond rotations in both conformer **1** and conformer **2**, such as the C17‒ C18/C17’‒C18’ and C18‒C19/C18’‒C19’ bonds (Figures S5 and S6 in Supporting Information). Other rotations, such as the C4‒C7/C4’‒C7’ (phenylethanolamine group) and C14‒N2/C14’‒N2’ (acetanilide group) bonds, were necessary to better accommodate the chemical environment within the protein binding pocket and involved rotations exceeding 50° (Figures S5 and S6 in Supporting Information).

**Figure 3.**
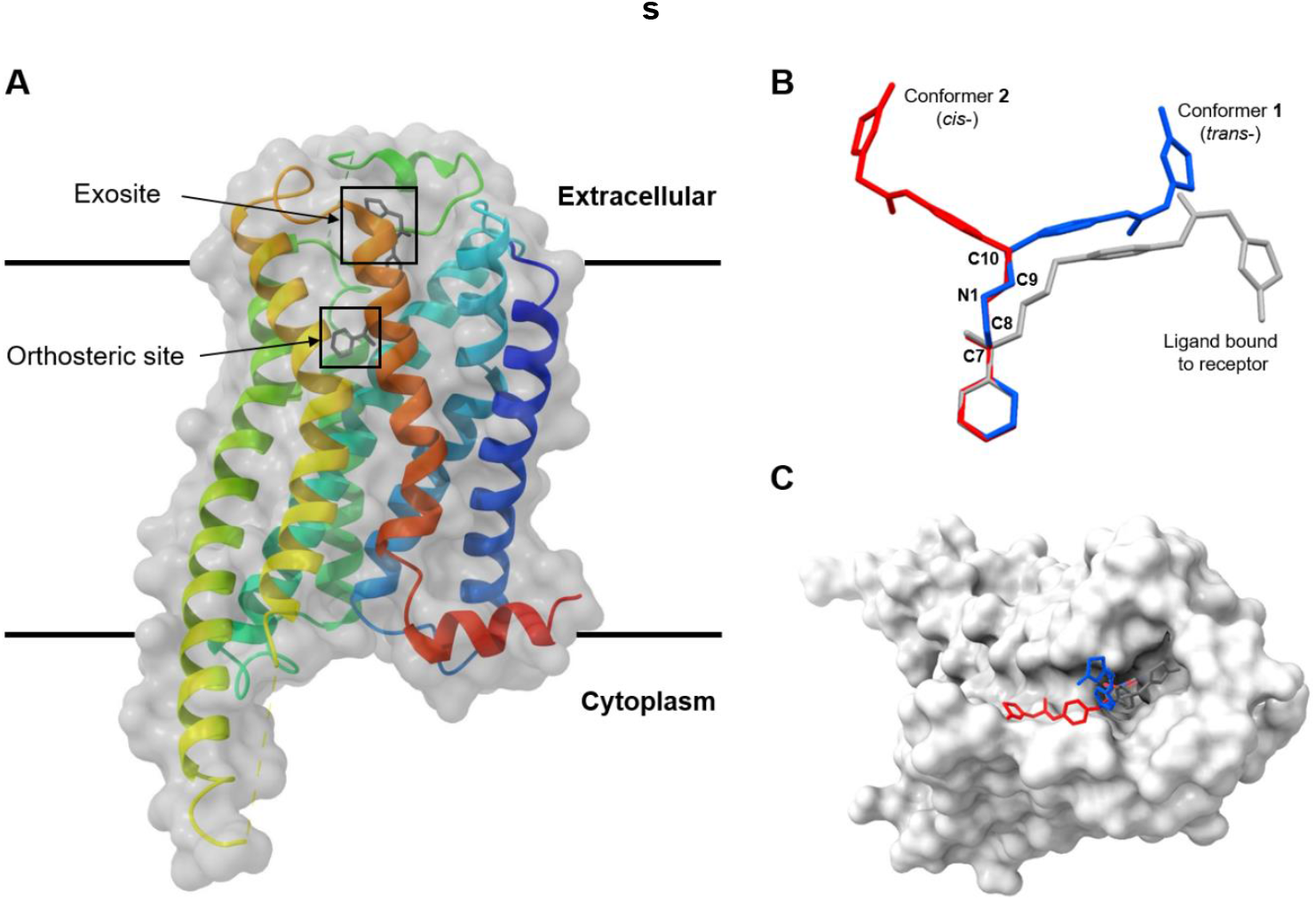
Overlay of MicroED and Cryo-EM structures of Mirabegron inside of the dog β3AR (PDB entry: 7DH5).^1^ (A) Overall view; (B) Overlay of conformer **1** and **2** (MicroED) and Cryo-EM structure; (C) View of the Mirabegron binding pocket viewed from the extracellular side of the membrane. Conformer **1** was colored in blue, conformer **2** was colored in red, Cryo-EM ligand structure was colored in grey, β3AR was colored in white surface. H atoms were omitted for clarity. This analysis shows that Mirabegron undergoes a large conformational change upon binding to the receptor as the structures of the drug alone (conformers **1** and **2**) cannot fit otherwise.

In summary, the atomic resolution structure of Mirabegron was successfully determined using MicroED, utilizing nanocrystals directly from seemingly amorphous powder. The structure revealed the presence of two distinct conformers within the lattice, packed in a “head-to-tail” arrangement. The hydrophilic groups of the molecule were embedded within the pairs, resulting in a hydrophobic surface and low water solubility. Comparison of the structures showed the presence of *trans*- and *cis*-forms, primarily determined by the rotation of C9‒C10/C9’‒C10’ bonds. Neither conformer of the drug alone matches the structure of Mirabegron bound to its receptor as determined by cryoEM suggesting that the drug must undergo a significant conformational change to bind in the agonist binding pocket of β3AR. Alternatively, since the single particle cryoEM density for the drug was poor it is entirely possible that the binding site may be different, possibly the allosteric site that fits conformer 2 is a likely alternative. This study highlights the potential of MicroED for solving unknown and polymorphic structures of drugs to aid in precision drug docking and the design of drugs that undergo conformational changes for increased specificity.

## Supplementary information

Text, experimental details and additional data, including Scheme S1, Figures S1−S7 and Tables S1-S3.

## Acknowledgements

The authors thank Michael W. Martynowycz for support and discussions. This study was funded in part by the National Institutes of Health P41GM136508. Portions of this research or manuscript completion were developed with funding from the Department of Defense grants MCDC-2202-002. Effort sponsored by the U.S. Government under Other Transaction number W15QKN-16-9-1002 between the MCDC, and the Government. The US Government is authorized to reproduce and distribute reprints for Governmental purposes, notwithstanding any copyright notation thereon. The views and conclusions contained herein are those of the authors and should not be interpreted as necessarily representing the official policies or endorsements, either expressed or implied, of the U.S. Government. The PAH shall flowdown these requirements to its subawardees, at all tiers. The Gonen laboratory is supported by funds from the Howard Hughes Medical Institute.

## Reference

1 C. Nagiri, K. Kobayashi, A. Tomita, M. Kato, K. Kobayashi, K. Yamashita, T. Nishizawa, A. Inoue, W. Shihoya, O. Nureki, Mol. Cell., 2021, 81, 3205–3215.

2 E. Sacco, R. Bientinesi, D. Tienforti, M. Racioppi, G. Gulino, D. D’Agostino, M. Vittori, P. Bassi, Expert Opin. Drug Discov., 2014, 9, 433–448.

3 T. Takasu, M. Ukai, S. Sato, T. Matsui, I. Nagase, T. Maruyama, M. Sasamata, K. Miyata, H. Uchida, O. Yamaguchi, J. Pharmacol. Exp. Ther., 2007, 321, 642–647.

4 J.-H. An, C. Lim, A. N. Kiyonga, I. H. Chung, I. K. Lee, K. Mo, M. Park, W. Youn, W. R. Choi, Y.-G. Suh, Pharmaceutics., 2018, 10, 149.

5 J. M. Belavic, Nurse. Pract., 2013, 38, 24–42.

6 S. Kane, ClinCalc DrugStats Database, 2020, 20.

7 J. H. Q. Mendoza, J. A. Henao, A. P. Aparicio, A. R. R. Bohorquez, Powder Diffr., 2017, 32, 290–294.

8 A. P. Tamayo, R. A. Toro, J. A. Henao, Rev. Invest. Univ. Quindío., 2019, 31.

9 K. Diederichs, M. Wang, Protein Crystallography: Methods and Protocols 2017, 239–272.

10 T. K. Brunck, F. Weinhold, J. Am. Chem. Soc., 1979, 101, 1700–1709.

11 D. Shi, B. L. Nannenga, M. G. Iadanza, T. Gonen, elife, 2013, 2, e01345.

12 B. L. Nannenga, D. Shi, A. G. W. Leslie, T. Gonen, Nat. Methods., 2014, 11, 927–930.

13 Y. Wang, S. Takki, O. Cheung, H. Xu, W. Wan, L. Öhrström, A. K. Inge, ChemComm, 2017, 53, 7018–7021.

14 T. Gruene, J. T. C. Wennmacher, C. Zaubitzer, J. J. Holstein, J. Heidler, A. Fecteau-Lefebvre, S. De Carlo, E. Müller, K. N. Goldie, I. Regeni, Angew. Chem. Int. Ed., 2018, 57, 16313–16317.

15 I. Andrusenko, V. Hamilton, E. Mugnaioli, A. Lanza, C. Hall, J. Potticary, S. R. Hall, M. Gemmi, Angew. Chem. Int. Ed., 2019, 58, 10919–10922.

16 L. J. Kim, M. Xue, X. Li, Z. Xu, E. Paulson, B. Mercado, H. M. Nelson, S. B. Herzon, J. Am. Chem. Soc., 2021, 143, 6578–6585.

17 D. P. Karothu, Z. Alhaddad, C. R. Göb, C. J. Schürmann, R. Bücker, P. Naumov, Angew. Chem. Int. Ed., 2023, e202303761.

18 S. Sekharan, X. Liu, Z. Yang, X. Liu, L. Deng, S. Ruan, Y. Abramov, G. Sun, S. Li, T. Zhou, RSC Adv., 2021, 11, 17408–17412.

19 M. Lightowler, S. Li, X. Ou, X. Zou, M. Lu, H. Xu, Angew. Chem. Int. Ed., 2022, 61, e202114985.

20 S. Li, M. Lightowler, X. Ou, S. Huang, Y. Jiang, X. Li, X. Zou, H. Xu, M. Lu, Commun. Chem., 2023, 6, 18.

21 T. Sasaki, T. Nakane, A. Kawamoto, T. Nishizawa, G. Kurisu, CrystEngComm, 2023.

22 D. Gogoi, T. Sasaki, T. Nakane, A. Kawamoto, H. Hojo, G. Kurisu, R. Thakuria, ChemRxiv, 2023.

23 J. E. Burch, A. G. Smith, S. Caille, S. D. Walker, R. Wurz, V. J. Cee, J. Rodriguez, D. Gostovic, K. Quasdorf, H. M. Nelson, ChemRxiv, 2021.

24 C. G. Jones, M. W. Martynowycz, J. Hattne, T. J. Fulton, B. M. Stoltz, J. A. Rodriguez, H. M. Nelson, T. Gonen, ACS Cent. Sci., 2018, 4, 1587–1592.

25 J. Hattne, M. W. Martynowycz, P. A. Penczek, T. Gonen, IUCrJ, 2019, 6, 921–926.

26 W. Kabsch, Acta Crystallogr., Sect D: Biol. Crystallogr., 2010, 66, 125–132.

27 W. Kabsch, Acta Crystallogr., Sect D: Biol. Crystallogr., 2010, 66, 133–144.

28 G. M. Sheldrick, Acta Crystallogr., Sect. A: Found. Crystallogr. Advances., 2015, 71, 3–8.

29 G. M. Sheldrick, Acta Crystallogr., Sect. C: Cryst. Struct. Commun., 2015, 71, 3–8.

30 S. Tsuzuki, Intermolecular forces and clusters I, 2005, 149-193.

31 J. Garbarczyk, G. Kamyszek, R. Boese, J. Mol. Struct. 1999, 479, 21–30.

32 Crystallographic Information File (CIF) of Mirabegron is deposited in Cambridge Crystallographic Data Center (CCDC) and the number is 2277685.

