## Supplemental Information for "Distinct Conformations of Mirabegron Determined by MicroED"

### Methods

#### Materials.

Mirabegron (2-(2-Amino-1,3-thiazol-4-yl)-*N*-[4-(2-{[(2*R*)-2-hydroxy-2-phenylethyl]amino}ethyl)phenyl]acetamide) was commercially purchased from InvivoChem and used as received without further recrystallization.

#### Grid preparation.

Sample preparation followed procedure as described previously.[1] One carbon-coated copper grid (400-mesh, 3.05 mm O.D., Ted Pella Inc.) was pretreated with glow-discharge plasma at 15 mA on the negative mode using PELCO easiGlow (Ted Pella Inc.) for 60s. Around 1 mg of powdery compounds were carefully weighed by a Mettler Toledo (XPR225DR) analytical balance and mixed with a grid in a 10 mL scintillation vial. After gently shaking the vial, the grid was removed and clipped at room temperature.

#### MicroED data collection.

The clipped grid was loaded in an aligned Thermo Fisher Talos Arctica Cryo-TEM (200 kV, ~0.0251 Å) at 100 K, equipped with a CetaD CMOS camera (4096 × 4096 pixels) and EPUD (Thermo Fisher) software.[1,2] Screening of size- and thickness-suitable microcrystals was done in the imaging mode (LM 210× and SA 3400×). The MicroED data was collected in the diffraction mode with 741 mm diffraction length, 70 µm C2 aperture, and a 50 µm selected area (SA) aperture in the parallel beam condition (45.2% C2 intensity) which resulted in a beam size at approximately 1.4 µm. Typical data collection used a constant rotation rate of ~1° per second over an angular wedge of 100° or 120° from -50° to +50° or -60° to +60°, respectively, with 1s exposure time per frame. Crystals selected for MicroED data collection were isolated and calibrated to eucentric height to maintain the crystal inside the beam during the rotation.

**MicroED data processing.**

The MicroED data was saved in mrc format and converted to smv format using the mrc2smv software (https://cryoem.ucla.edu/microed).[2] The converted frames were indexed and integrated by XDS.[3,4] Three selected datasets with the highest resolution at 0.9 Å were scaled and merged using XSCALE,[4] and intensities were converted to SHELX hkl format using XDSCONV.[4] The merged dataset showed 99.7% overall completeness, which can be *ab initio* solved by SHELXT[5] with a resolution of 1.01 Å. The structure was refined by SHELXL[6] in Shelxle[7] as a graphical interference to yield the final MicroED structure (Figure 1, Table S1 in Supporting Information).

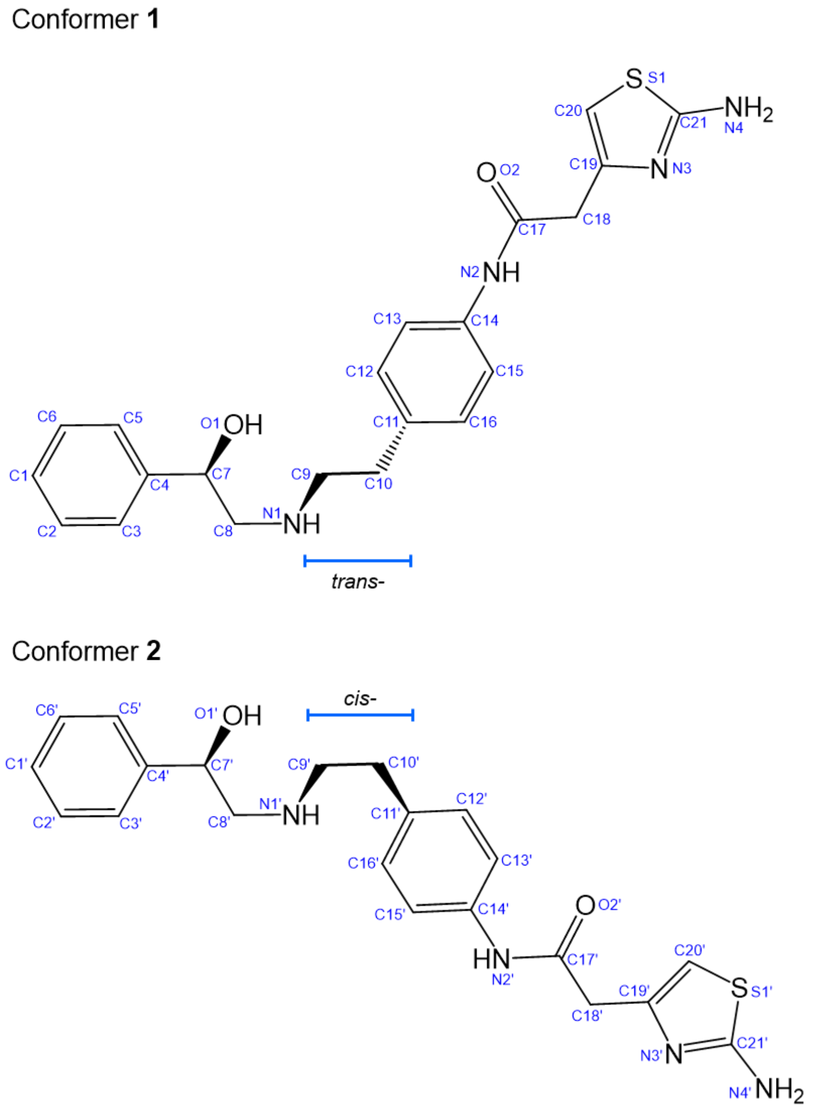


**Scheme S1** Chemical notations of Mirabegron. Conformer **1** was labeled with atom type and numbers, conformer **2** was labeled with atom type and primed numbers. Conformations along C9–C10 and C9'–C10' were highlighted, showing the *trans*- and *cis*- form in conformer **1** and **2**, respectively.

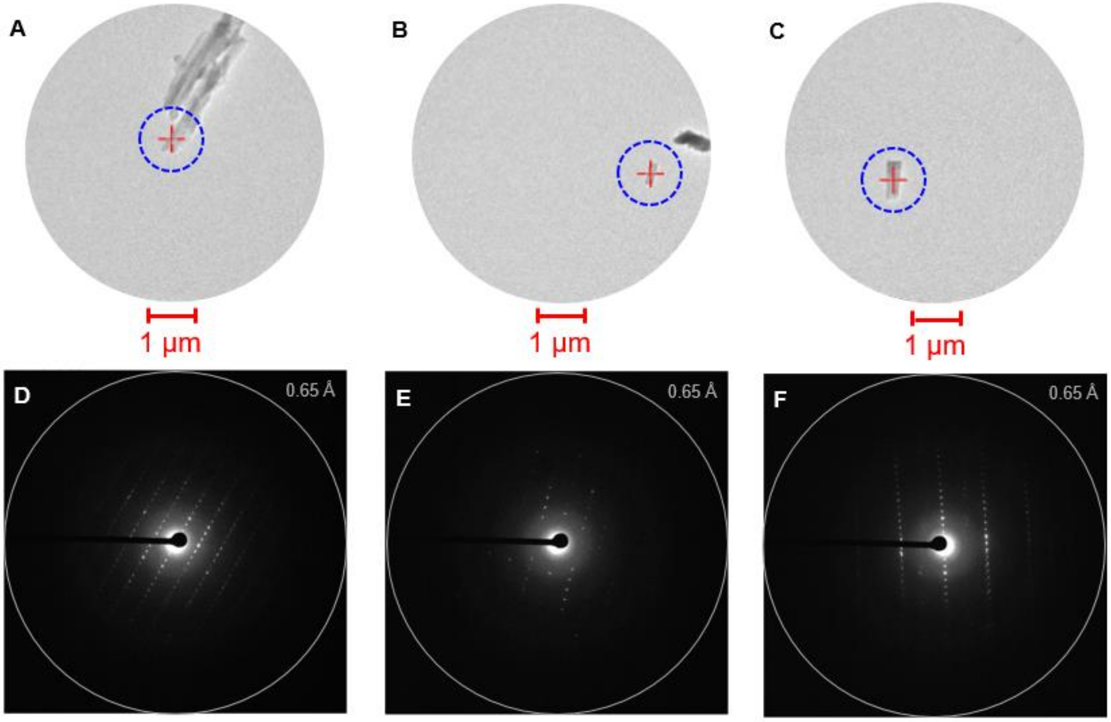


**Figure S1** Crystal appearance and diffraction pattern under the TEM. (A-C) Images of items 1-3 under imaging mode (SA 3400×), the diffraction beam size was highlighted in dashed blue circles; (D-F) Diffraction pattern of items 1-3 under diffraction mode (741 mm), the integration edge was colored in grey rings.

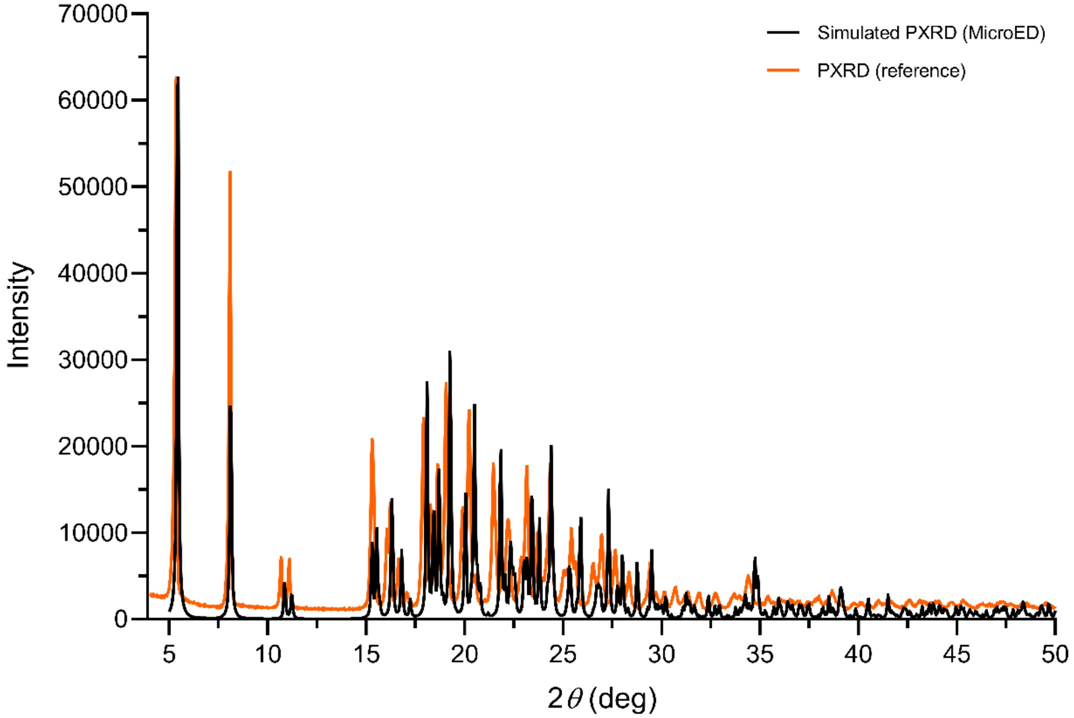


**Figure S2** Overlay of literature-reported and simulated PXRD spectra of Mirabegron.[8] Literature-reported PXRD data was colored in orange line; simulated PXRD data was back-calculated from MicroED structure and colored in black line. Intensities were rescaled for comparison.

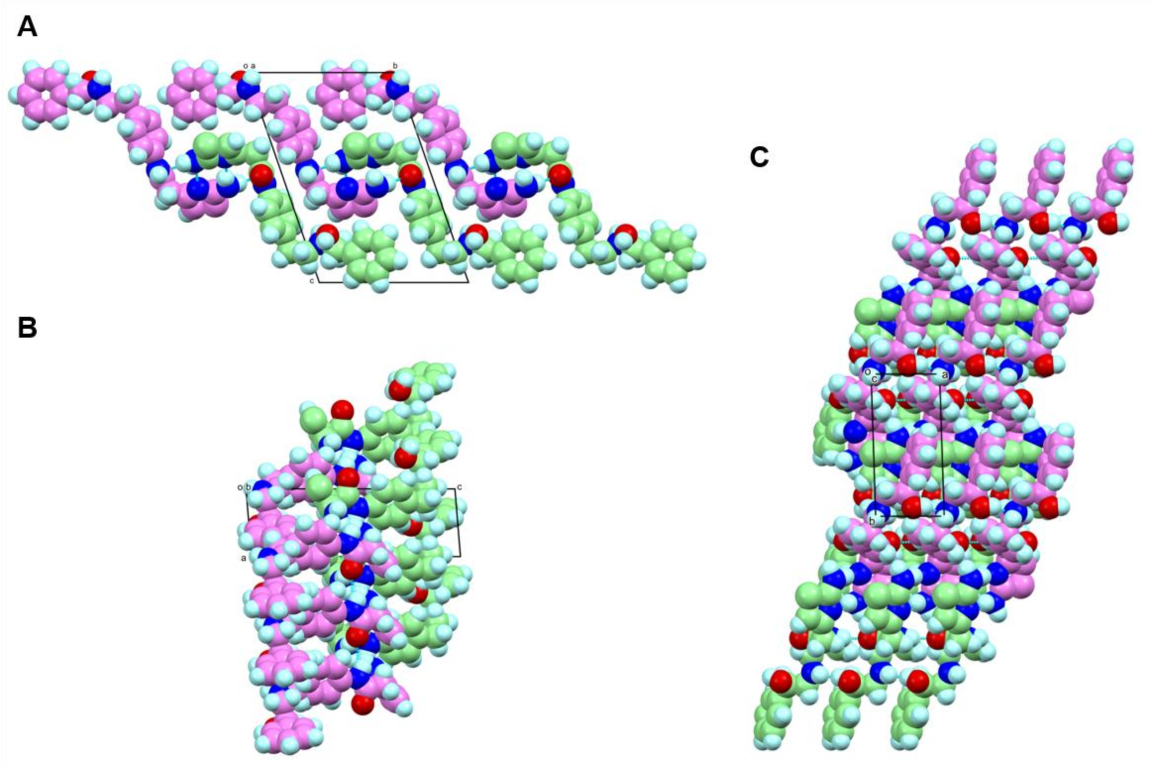


**Figure S3** Dense packing of Mirabegron observed in the crystal lattice. (A) viewed along *a* axis; (B) viewed along *b* axis; (C) viewed along *c* axis. Conformer **1** was colored in violet, conformer **2** was colored in light green. The cyan dashed lines represented the hydrogen-bond interactions, with the contact oxygen and nitrogen atoms colored in red and blue, respectively. The hydrogen atoms were colored in cyan.

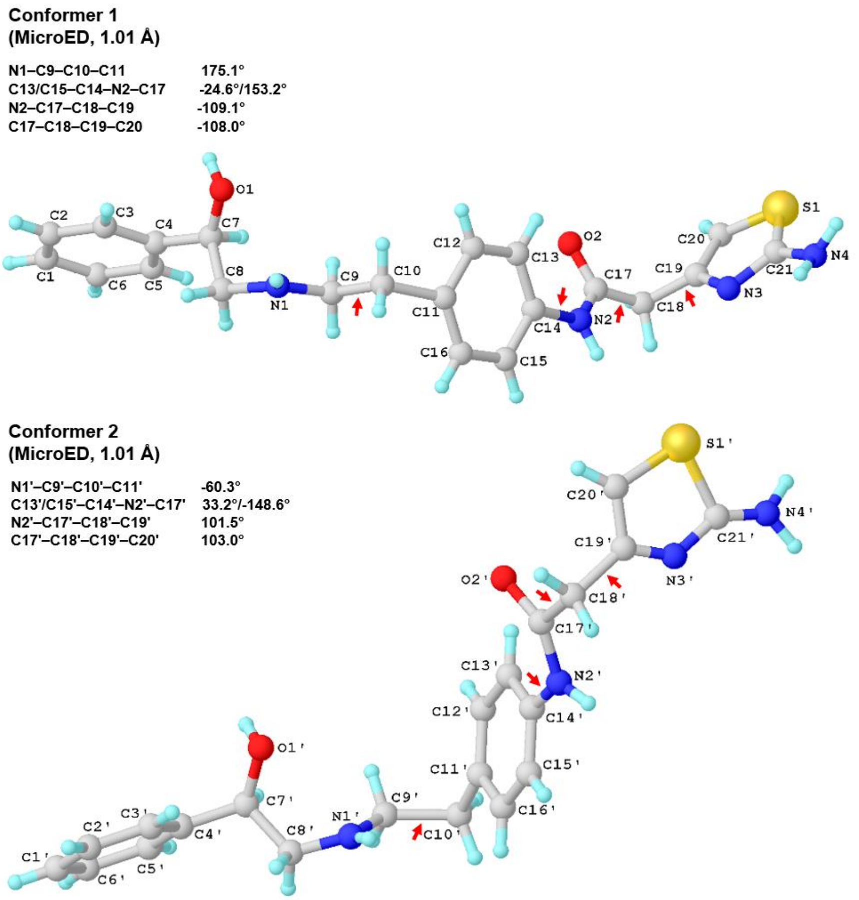


**Figure S4** Major structural differences observed in conformer **1** and **2**. The primary torsion differences were highlighted by red arrows. Selected torsion angles were listed for comparison.

**Conformer 1**
**(MicroED, 1.01 Å)**

| | |
|---|---|
| C3/C5–C4–C7–C8 | 84.4°/-92.3° |
| C7–C8–N1–C9 | -74.1° |
| C13/C15–C14–N2–C17 | -24.6°/153.2° |
| N2–C17–C18–C19 | -109.1° |
| C17–C18–C19–C20 | -108.0° |

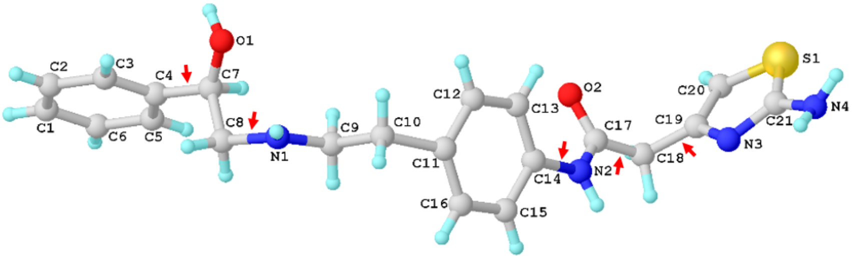


**Ligand**
**(CryoEM, 3.16 Å)**

| | |
|---|---|
| C3/C5–C4–C7–C8 | 147.9°/-32.0° |
| C7–C8–N1–C9 | 167.3° |
| C13/C15–C14–N2–C17 | -103.8°/76.8° |
| N2–C17–C18–C19 | 32.2° |
| C17–C18–C19–C20 | 123.7° |

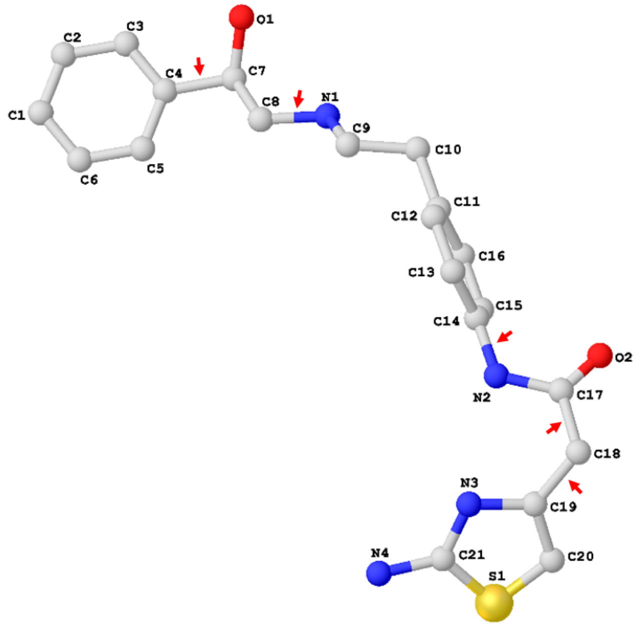


**Figure S5** Major structural differences observed in conformer **1** and Cryo-EM structure.[9] The primary torsion differences were highlighted by red arrows. Selected torsion angles were listed for comparison. H atoms were omitted for clarity.

**Conformer 2**
**(MicroED, 1.01 Å)**

| | |
|---|---|
| C3'/C5'–C4'–C7'–C8' | 78.6°/-100.8° |
| C7'–C8'–N1'–C9' | -59.3° |
| N1'–C9'–C10'–C11' | -60.3° |
| C13'/C15'–C4'–N2'–C17' | 33.2°/-148.6° |
| N2'–C17'–C18'–C19' | 101.5° |

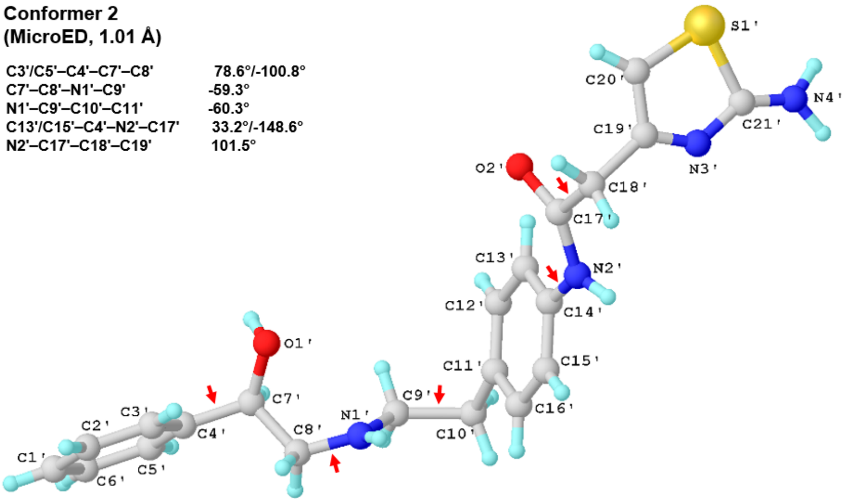


**Ligand**
**(CryoEM, 3.16 Å)**

| | |
|---|---|
| C3/C5–C4–C7–C8 | 148.9°/-32.0° |
| C7–C8–N1–C9 | 167.3° |
| N1–C9–C10–C11 | -143.0° |
| C13/C15–C4–N2–C17 | -103.8°/76.8° |
| N2–C17–C18–C19 | 32.3° |

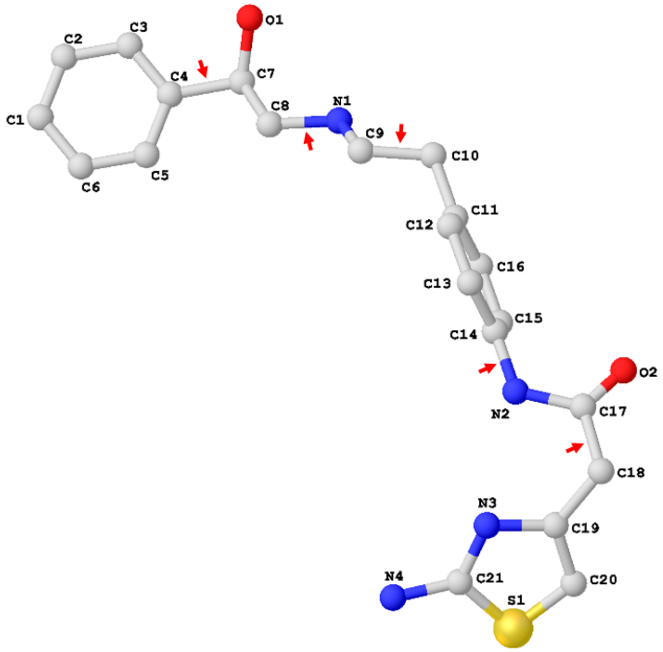


**Figure S6** Major structural differences observed in conformer **2** and Cryo-EM structure.[9] The primary torsion differences were highlighted by red arrows. Selected torsion angles were listed for comparison. H atoms were omitted for clarity.

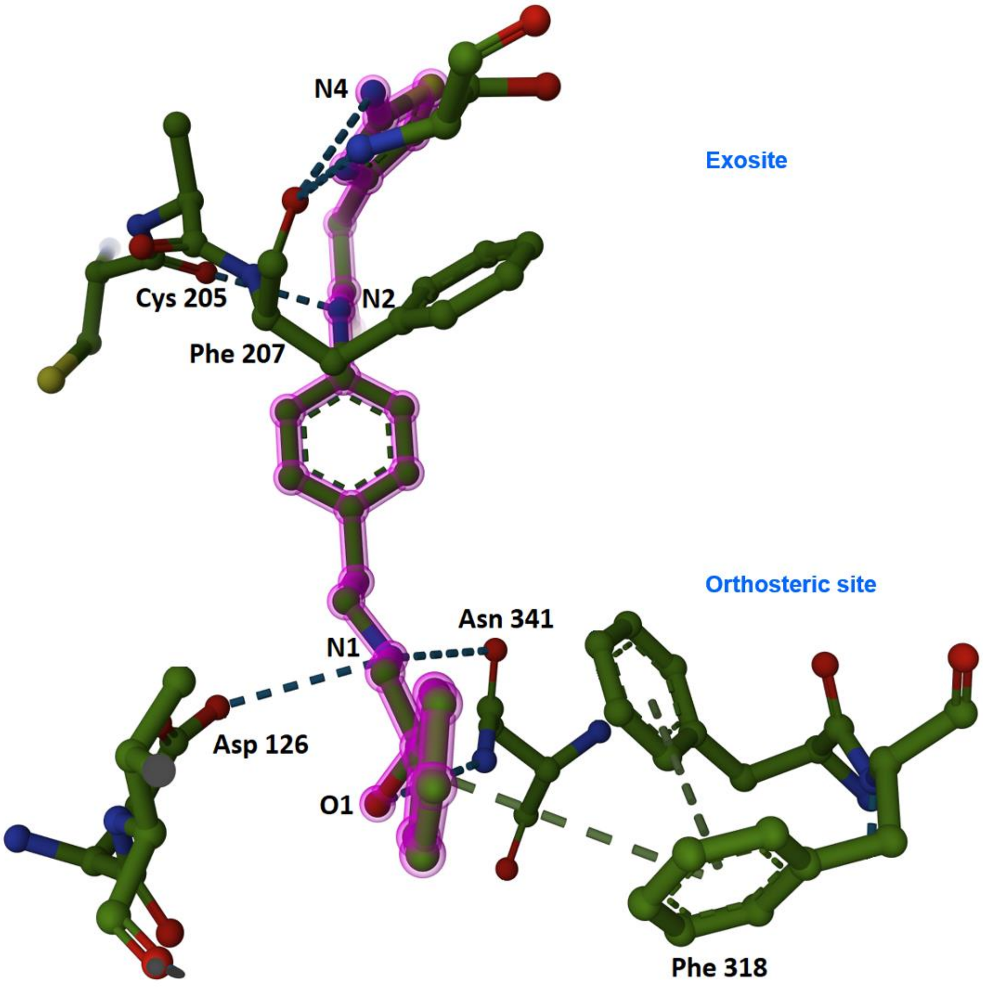


**Figure S7** Hydrogen bonding and *van der Waals* interactions between Mirabegron and the active sites of β3AR (PDB entry: 7DH5).[9] Mirabegron was highlighted in violet, and the contact atoms and the residues involved in hydrogen bonding were labeled. H atoms were omitted for clarity.

**Table S1** MicroED data statistics of Mirabegron (merged).

| | |
|---|---|
| Stoichiometric formula | $C_{21}H_{24}N_4O_2S$ |
| Mr | 396.50 |
| Temperature (K) | 100 |
| Crystal system | Triclinic |
| Space group | P1 |
| Unit cell lengths (Å) | |
| a | 5.27 |
| b | 11.58 |
| c | 17.27 |
| Unit cell angles (°) | |
| α | 70.731 |
| β | 84.351 |
| γ | 86.372 |
| Cell volume ($Å^3$) | 998.51 |
| No. of observed reflections | 7518 |
| No. of unique reflections | 4014 |
| $R_{obs}$ (%) | 18.2 |
| $R_{meas}$ (%) | 24.8 |
| I/Sigma | 3.09 |
| $CC_{1/2}$ | 95.5 |
| Resolution (Å) | 1.01 |
| Completeness (%) | **99.7** |
| $R_1$ (%) | **16.59** |
| $wR_2$ (%) | 40.17 |
| GooF | 1.288 |

**Table S2** MicroED data statistics of three selected items of Mirabegron.

| | Item 1 | Item 2 | Item 3 |
|---|---|---|---|
| Space group | P1 | P1 | P1 |
| Unit cell lengths (Å) | | | |
| a | 5.27 | 5.28 | 5.27 |
| b | 11.58 | 11.41 | 11.66 |
| c | 17.27 | 17.37 | 17.18 |
| Unit cell angles (°) | | | |
| α | 70.731 | 70.441 | 71.265 |
| β | 84.351 | 84.632 | 84.377 |
| γ | 86.372 | 86.375 | 86.254 |
| No. of observed reflections | 6878 | 6314 | 4856 |
| No. of unique reflections | 3547 | 3245 | 2682 |
| $R_{obs}$ (%) | 18.3 | 18.9 | 14.4 |
| $R_{meas}$ (%) | 25.9 | 26.7 | 20.4 |
| I/SIGMA | 2.39 | 3.23 | 3.12 |
| $CC_{1/2}$ | 97.9 | 98.4 | 98.6 |
| Resolution (Å) | 0.9 | 0.9 | 0.9 |
| Completeness (%) | 72.2 | 66.7 | 54.2 |

**Table S3** Hydrogen-bond geometry in Mirabegron (Å, °)

| **D–H···A** | **D–H** | **H···A** | **D···A** | **D–H···A** |
|---|---|---|---|---|
| O1–H···N1$^{i}$ | 0.825 | 2.060 | 2.797 | 148.59 |
| N2–H···O2$^{ii}$ | 0.865 | 2.514 | 3.275 | 147.28 |
| N4–H···O2'$^{i}$ | 0.895 | 2.017 | 2.907 | 172.35 |
| N4–H···N3' | 0.918 | 2.294 | 3.195 | 167.03 |
| O1'–H···N1'$^{ii}$ | 0.839 | 2.077 | 2.853 | 153.61 |
| N2'–H···O2'$^{i}$ | 0.867 | 2.117 | 2.969 | 167.58 |
| N4'–H···O2$^{ii}$ | 0.928 | 2.142 | 3.063 | 171.81 |
| N4'–H···N3 | 0.935 | 2.283 | 3.086 | 143.69 |
| Symmetry codes: *(i) x+1, y, z; (ii) x−1, y, z.* | | | | |

### Reference


1. C. G. Jones, M. W. Martynowycz, J. Hattne, T. J. Fulton, B. M. Stoltz, J. A. Rodriguez, H. M. Nelson, T. Gonen, *ACS Cent. Sci.,* **2018**, *4*, 1587-1592.
2. J. Hattne, M. W. Martynowycz, P. A. Penczek, T. Gonen, *IUCrJ,* **2019**, *6*, 921-926.
3. W. Kabsch, *Acta Crystallogr., Sect D: Biol. Crystallogr.,* **2010**, *66*, 125-132.
4. W. Kabsch, *Acta Crystallogr., Sect D: Biol. Crystallogr.,* **2010**, *66*, 133-144.
5. G. M. Sheldrick, *Acta Crystallogr., Sect. A: Found. Crystallogr. Advances.,* **2015**, *71*, 3-8.
6. G. M. Sheldrick, *Acta Crystallogr., Sect. C: Cryst. Struct. Commun.,* **2015**, *71*, 3-8.
7. C. B. Hübschle, G. M. Sheldrick, B. Dittrich, *J. Appl. Crystallogr.,* **2011**, *44*, 1281-1284.
8. J. H. Q. Mendoza, J. A. Henao, A. P. Aparicio, A. R. R. Bohorquez, *Powder Diffr.,* **2017**, *32*, 290-294.
9. C. Nagiri, K. Kobayashi, A. Tomita, M. Kato, K. Kobayashi, K. Yamashita, T. Nishizawa, A. Inoue, W. Shihoya, O. Nureki, *Mol. Cell.,* **2021**, *81*, 3205-3215.
